# Targeted Quantitative Analysis of Polar and Phosphorylated Intermediates of Algal and Plant Primary Carbon Metabolism Using an Optimized HILIC-Amide Tandem MS Workflow

**DOI:** 10.64898/2026.09.28.754878

**Authors:** Nguyen Kim Ngan Bui, David Achaintre, Frederik Sommer, Ayla Khold Atila, Or Geffen, Melissa Meinert, Gloria Okooboh, Haim Treves

## Abstract

Central carbon metabolism coordinates energy production and carbon skeleton allocation across photosynthetic and heterotrophic organisms. Here, we established an optimized targeted metabolomics workflow for *Chlamydomonas reinhardtii* and *Arabidopsis thaliana* using a single-run HILIC-MS/MS assay with alternating positive and negative ionization. Our method provides robust multi-class coverage across 21 amino acids, 11 carboxylic acids, 4 sugars, 3 adenine nucleotides, 2 nucleotide sugars, 18 phosphorylated compounds in CCM, and Acetyl-CoA in a 20-minute run using both ESI modes. An optimal concentration of Medronic acid with a BEH Amide column at high pH prevented metal adsorption, and low quantification limits (10 to 50 nM) for labile intermediates such as PEP, ATP and RuBP were achieved. Retention time variations remained below 1.3%, and over 90% of analytes showed high linearity (R2 > 0.990) across dynamic ranges spanning two to three orders of magnitude (typically 0.01-0.5 µM up to 15-20 µM).

For sample preparation, glass-fiber filters (GF/C) outperformed nitrocellulose for harvesting *C. reinhardtii*. For *A. thaliana*, it was proved that no further grinding with micro pestle and extended cooling were necessary. A two-phase chloroform/methanol/water extraction was essential, as a single-phase extraction co-extracted green chlorophyll, which heavily suppressed phosphorylated signals. In Arabidopsis, glutamate, fumaric acid and citric acid were the dominant primary pools, indicating the importance of dilution and column washing for preventing carryover, and sensitive intermediates were reliably quantified at physiological levels. Together, this unified analytical and extraction strategy enables robust, artifact-free, high-throughput profiling of central metabolism in photosynthetic organisms.

## 1. Introduction

Central carbon metabolism (CCM) provides the carbon skeletons and energy required for growth in plants and algae [1]. Photosynthetically fixed carbon enters the Calvin–Benson cycle and is distributed through interconnected pathways, including glycolysis [2], the pentose phosphate pathway, and the tricarboxylic acid cycle, to support the synthesis of sugars, organic acids, amino acids, and other cellular components [3,4]. Although these pathways are broadly shared, carbon partitioning differs among plant and algal species and in response to growth conditions, making their metabolites valuable targets for metabolic profiling [5,6].

Despite its biological importance, the quantitative profiling of CCM intermediates presents severe analytical bottlenecks. These analytes exhibit extreme hydrophilicity, high negative charge density, low thermal volatility, structural isomerism, and a strong propensity for non-specific metal chelation. Traditional gas chromatography-mass spectrometry (GC-MS) workflows require labor-intensive chemical derivatization [7,8]. Conversely, conventional reverse-phase liquid chromatography (RP-C18) fails to retain highly polar species, causing them to elute in void volume [9]. While RP ion-pairing chromatography (RP-IPC) utilizing volatile alkylamines (e.g., tributylamine) achieves retention of anionic sugar phosphates, the millimolar additive concentrations cause severe electrospray ionization (ESI) suppression and persistent LC-MS system contamination [10,11]. Alternative approaches such as capillary electrophoresis (CE-MS) and anion-exchange chromatography offer high selectivity for polyanions but often suffer from lower chromatographic robustness or non-volatile salt gradients that foul ESI sources [12,13].

Hydrophilic interaction liquid chromatography (HILIC) has emerged as the premier platform for direct polar metabolome profiling without derivatization or ion-pairing agent [14]. Early untargeted HILIC screening frequently employed zwitterionic stationary phases (e.g., SeQuant ZIC-pHILIC) under neutral or weakly basic conditions, providing broad polarity coverage [15]. However, recent systematic evaluations comparing bioinert stationary phases have demonstrated that amide-linked hybrid silica phases (such as Acquity Premier BEH Amide) offer superior chromatographic efficiency, narrower peak widths, and higher retention time repeatability over bare silica and zwitterionic phases [16]. Furthermore, operating HILIC separations at basic pH (pH 9.0–11.0) maximizes negative ESI ionization and dramatically accelerates the mutarotation kinetics of reducing sugar phosphates, effectively collapsing isomeric anomeric doublets into single sharp peaks [13,16,17].

A major remaining challenge in HILIC-MS/MS is the non-specific adsorption of polyphosphorylated analytes (e.g., ATP, RuBP, FBP) to active metal sites on stainless-steel tubing, column frits, and ESI hardware. To prevent peak tailing and signal loss caused by interactions between trace metals with phosphorylated compounds, three additives are commonly used: citrate [18], EDTA, pyrophosphoric acid, and medronic acid. While EDTA works well during sample preparation [19], adding it to mobile phases causes severe signal suppression and system contamination [20]. Pyrophosphoric acid improves peak shapes, but it quickly breaks down in water, making long runs unreliable. In contrast, a standard 5 µM concentration medronic acid [21–23] usually serves as the benchmark baseline for general LC deactivation without inducing electrospray ionization (ESI) suppression. While lower micro-molar levels (2.5 – 7 µM) are tailored for targeted profiling of carboxylates, nucleotides, and isotopic tracer experiments [24]. Higher micro-molar and percentage additives (20 µM or 0.1% w/v) are applied to improve peak shape and chromatographic recovery for polyphosphates, purines, and antiviral drugs [25,26].

In this work, we developed a complementary, robust HILIC-MS/MS method by supplementing aqueous mobile phase A with an optimized concentration of medronic acid, while maintaining a low-salt regime (10 mM buffer, pH 9.0) at moderate column temperatures. After validation, this approach was applied along with optimized extraction protocols, to reliably measure nearly 60 central carbon metabolites in *Chlamydomonas reinhardtii* and *Arabidopsis thaliana*.

## 2. Methods and materials

Formic acid, ammonium hydroxide, ammonium acetate and medronic acid were purchased from Sigma-Aldrich. Acetonitrile, methanol and isopropanol of Ultra LC-MS grade were purchased from VWR. Ultra-pure water was obtained using a Milli-Q purification system. All metabolite standards (Table S1) were purchased from Merck (encompassing Sigma, Aldrich and Supelco product lines), with the exception of specific compounds obtained from other suppliers: BLD Pharmatech (Isocitric acid trisodium), TCI (2-Oxoglutaric acid, Shikimic acid and Adenosine 5’-diphosphate disodium), Thermo Fisher Scientific (Adenosine 5’-monophosphate disodium), Santa Cruz Biotechnology / SCBT (Uridine diphosphate glucose disodium, 2-Phosphoglyceric acid sodium, D-Erythrose 4-phosphate, D-Mannose 6-phosphate, and D-Ribulose 1,5-bisphosphate sodium), Chempur (D-Fructose 1,6-bisphosphate, Glucose 1-phosphate disodium, and D-Xylulose 5-phosphate), and BioSynth (D-Ribulose 5-phosphate and D-Sedoheptulose-7-phosphate barium). Ribitol was used as internal standard.

### 2.1. Cultivation and growth conditions

#### Microalgal Cultivation

***Chlamydomonas reinhardtii*** strain CC-5325 was grown axenically in flat-panel glass photobioreactors (FMT-150, PSI) containing 1 L of sterile HP (HEPES-phosphate) medium adjusted to pH 7.2. Cultures were initiated at an optical density of OD_735_ = 0.02 and maintained under continuous illumination of 100 *μ*mol photons ⋅ m^−2^ ⋅ s^−1^ (100 *μ*E) provided by cool-white LED panels. The culture temperature was maintained constantly at 25°C. Ambient synthetic air containing 0.04% CO_2_ (400 ppm) was bubbled continuously through the culture at a flow rate of approximately 1 L/min to supply inorganic carbon and ensure mixing. Culture growth and physiological parameters (pH, dissolved O_2_, and optical density at 680 and 735 nm) were monitored continuously using integrated bioreactor sensors. Triplicate biological replicates were cultivated independently for all experimental setups.

The rapid and slow sampling procedures differ primarily in the point of collection and the method of metabolic quenching [27,28]. In the rapid regime, external separate syringes for cell and 0.04% CO_2_ media were connected to the microfluidic mixing setup immediately deliver the mixed culture into chilled −70°C/ 70% methanol to arrest enzymatic activity within seconds, targeting highly dynamic turnover intermediates. In contrast, the slow regime samples live cell suspensions directly from the photobioreactor vessels for collection by filtration, focusing on stable downstream pools and macromolecular end-products. To demonstrate the quantification method, the slow sampling procedure was carried out for the baseline (*T*0), samples were withdrawn directly from the running photobioreactor cultures bubbled with 0.04% CO_2_ air prior to any gas switch.

#### Plant growth condition

***Arabidopsis thaliana*** ecotype Columbia (Col-*0*) were sown on standard soil (type ED73, Einheitserde Patzer; Sinntal-Altengronau, Germany) with 10% (*v*/*v*) sand, stratified at 4 °C for 48 h, and then grown under short-day regime (10 h light/14 h dark) at 60% relative humidity and 120 μmol m^−2^ s^−1^ light intensity (NS1 Valoya growth lights, Helsinki, Finland) at 21 °C, representing standard growth conditions.

### 2.2. Metabolite extraction and sample preparation

#### Optimal protocol for Chlamydomonas reinhardtii

*Chlamydomonas reinhardtii* cultures (5 mL) were rapidly harvested under growth light via vacuum filtration using either 1.2 µm pore size, ~260 µm thickness glass fiber (GF/C) or 1 µm pore size, 130 – 150 µm thickness nitrocellulose (NC) filters, both are from Cytiva. The cell-containing filter was quickly transferred to a 2 mL safety-cap vial, immediately quenched in liquid nitrogen, and stored at −80°C. For cell disruption, frozen filter samples were supplemented with approx. 20 µL glass beads (<106 µm), 105 µL chloroform, 525 µL precooled 70% (v/v) methanol, and 10 µL of 20 µM Ribitol. Cell lysis was accomplished through three successive freeze–thaw cycles alternating between liquid nitrogen immersion and thawing on ice with vigorous vortexing. The liquid–liquid extraction was initiated by adding 280 µL of ice-cold Milli-Q water to the lysate, vortexing, and centrifuging at 14,000 rpm for 5 min at 0°C. The upper polar phase (methanol/water) was collected into one of the two 2 mL tubes. The remaining nonpolar chloroform phase was re-extracted twice by sequentially adding 560 µL of ice-cold Milli-Q water, vortexing, and centrifuging at 14,000 rpm for 5 min at 4°C. The combined aqueous extracts (approx. 1,920 µL total) were split equally into two 2 mL screw-cap tubes (approx. 960 µL each). Subsequently, 280 µL of ice-cold Milli-Q water was added to each tube to lower the final methanol concentration below 15% (v/v) to prevent vacuum flashing during freeze-drying. The extracts were frozen at −80°C, transferred on dry ice to preserve the cold chain, and lyophilized to dryness overnight. The lyophilized extracts from both tubes were combined and reconstituted in 100 µL of Milli-Q water by thoroughly rinsing the tube walls and caps. Finally, the reconstituted sample was filtered through a 0.22 µm hydrophilic 96-well filter plate and transferred into a 300 µL LC autosampler vial insert prior to LC-MS/MS analysis.

#### Optimal protocol for Arabidopsis thaliana

***Arabidopsis thaliana*** was harvested under the light and immediately homogenized in liquid nitrogen using a porcelain mortar and pestle. Weigh 20 mg (± 1 mg) of the homogenized tissue were weighed into 1.5 mL screw-cap microcentrifuge tubes maintained on ice. Metabolite extraction was initiated by adding 250 µL of an ice-cold (−20°C) chloroform/methanol mixture (3:7, v/v), followed by the addition of 10 µL of 50 µM Ribitol. The mixture was briefly vortexed and kept on ice. The effect of pestle (for cell strainer) after adding solvents and extended cooling 2h at −20°C are omitted and discussed in section 3.5.

Phase separation was induced by adding 400 µL of ice-cold ultra-pure water, vortexing thoroughly, and centrifuge at 14,000 rpm or maximum speed for 5 min at 4°C. The upper aqueous phase (approx. 650 µL) was collected and transferred to a 5 mL screw-cap tube. The remaining chloroform phase was re-extracted twice by sequentially adding 400 µL of ice-cold MilliQ water, vortexing, and centrifuging at 14,000 rpm or maximum speed (5 min, 4°C) for each cycle. The resulting upper phases (approx. 400 µL each) were pooled with the initial extract into the corresponding 5 mL tube. The combined extracts were frozen at −80°C and transferred to a lyophilizer using dry ice or liquid nitrogen to maintain the frozen state. Lyophilization was performed overnight through the perforated 5 mL screw-cap tubes. The resulting dry residues were reconstituted in 250 µL of Milli-Q water, ensuring complete rinsing of the tube walls to guarantee homogeneity. The reconstituted extracts were filtered through a Multiscreen 96-well filter plate (0.22 µm hydrophilic membrane, Millipore) without prior membrane wetting. Finally, 100 µL aliquots of the filtered samples were transferred into two separate HPLC vial inserts (one for primary LC-MS analysis and one as a backup archive) and stored at −80°C until analysis.

### 2.3. UHPLC-MS/MS

Targeted metabolite separation was conducted on an ACQUITY UPLC I-Class SM-FTN Bio system (Waters) composed of a binary pump, a flow-through-needle (FTN) autosampler and a heated column compartment, equipped with an Acquity Premier BEH Amide column (100 × 2.1 mm, 1.7 µm particle size; Waters) fitted with an Acquity Premier BEH Amide guard column. The mobile phase system consisted of 75 µM medronic acid (MDP) in Milli Q water (pH 4.0; mobile phase A) and 90% (v/v) acetonitrile containing 10 mM ammonium acetate (pH 9.0 adjust with ammonium hydroxide; mobile phase B). All mobile phases were stored in inert fluoropolymer bottles (Nalgene FEP) to mitigate effect of trace metal [29]. Chromatographic elution was executed at a flow rate of 0.4 mL/min with a column temperature maintained at 30°C, using seal wash (10% isopropanol) and needle wash solutions (acetonitrile/water, 80:20, v/v). The injection volume was 1 µL. The gradient program was operated as follows: 0 – 1.0 min, 90% B; 1.0 – 12.0 min, 90% – 75% B; 12.0 – 16.0 min, 75% – 50% B; 16.0 – 16.1 min, 50% – 90% B; and 16.1–20.0 min, re-equilibration at 90% B.

Mass spectrometric detection was performed on Sciex QTRAP 6500+ operating in multiple reaction monitoring (MRM) mode with electrospray ionization (ESI), software Sciex OS 3.4.5. The Turbo V ion source optimized parameters were as follows: curtain gas 35 psi, ion source gas 1 (spray gas) 50 psi and gas 2 (auxiliary gas) 60 psi, CAD gas 9, ion source spray voltage 5500 V for positive mode and −4500 V for negative mode, Q1 and Q3 High resolution and source temperature 400°C for all metabolites except Fructose and Glucose (200°C). Detail metabolite MRM transitions optimal parameters are summarized in Table S2 and S3.

### 2.4. Method validation

#### Preparation of Calibration Standards

Individual metabolite stock solutions were prepared by dissolving appropriate amounts of each standard in 30% methanol, with the exceptions of asparagine, glutamine, tyrosine, and cystine, which were dissolved in 0.1 M HCl. An intermediate mixture containing all metabolites at a concentration of approximately 300 μM was prepared in 30% methanol. This intermediate solution was then serially diluted to yield calibration standards ranging from 5 nM to 20 μM for each metabolite containing 2 μM of Ribitol. All calibrants were stored at −80 °C and thawed only once for single use. LOD and LOQ for each metabolite were determined by signal/noise (S/N) ratio equals to 3 and 10 respectively. Weighted least-squares regression (1/x) was applied for all cases.

## 3. Results and discussion

### 3.1. LC condition

Although Hosseinkhani et al. [15] recommended SeQuant ZIC-pHILIC over BEH Amide column for global plasma metabolomics, their detailed chromatographic profiles revealed severe peak broadening for hexose monophosphates under their optimized conditions. In contrast, operating the BEH Amide column at alkaline pH (pH 9.0) provides superior peak sharpness and accelerates anomeric mutarotation for monophosphates, an advantage strongly supported by Langová et al. [16].

While the authors also suggested that metal-chelating additives such as medronic acid are redundant for monophosphorylated sugars and nucleosides using bioinert UHPLC systems. Our evaluations demonstrated that sugar bisphosphates (FBP and RuBP) behave differently. Due to the presence of two localized, highly charged phosphate moieties on a flexible carbohydrate backbone, these bisphosphates exhibit strong chelation potential. On the BEH Amide stationary phase, the addition of low concentrations of medronic acid (MDP) improved peak symmetry, eliminated peak tailing and enhanced signal-to-noise ratios for phosphorylated compounds.

As shown in Figure S1, increasing the MDP concentration from 10 µM (blue) to 20 µM (magenta) substantially enhanced peak intensity, improved peak symmetry, and eliminated tailing across both sugar bisphosphates and adenine nucleotides. In contrast, the addition of 0.01% formic acid to the 20 µM MDP condition (orange) delayed retention times and caused severe signal suppression (reducing peak heights by over two-fold), likely because acidification disrupted optimal basic HILIC retention mechanisms and negative-mode ionization efficiency.

To explore the upper threshold of passivation, higher MDP concentrations of 50 µM (red/blue) and 75 µM (green) were evaluated (Figure S2). While 50 µM MDP was sufficient to retain peak symmetry for most analytes, increasing the concentration to 75 µM MDP produced sharper, more symmetrical peak profiles, amplified peak intensities for adenylates (AMP, ADP, ATP) and nucleotide sugars (ADP-Glc, UDP-Glc). Consequently, an alkaline mobile phase containing 75 µM MDP without acidic modifiers provided the highest sensitivity and sharpest peak shapes.

In our optimized HILIC-MS/MS workflow, aqueous mobile phase A is prepared with a nominal concentration of 75 µM medronic acid. However, because chromatographic separation and analyte elution occur across a 10% to 50% mobile phase A gradient, the maximum effective on-column concentration delivered to the stationary phase and ESI source at any point during analyte elution reaches a peak of 37.5 µM (at 50% A), with an initial baseline concentration of 7.5 µM (at 10% A).

As established by Langová et al. [16], basic amino acid as histidine achieves its best peak shape at pH 3.5, while pH 9.0 is optimal for glucose 6-phosphate (G6P), acetyl-CoA, and lactose. In our workflow, histidine performance was intentionally compromised in favor of the pH 9.0 regime, representing an acceptable trade-off to secure comprehensive, multi-class coverage in a single run to ensure superior peak symmetry and enhanced negative-mode ionization.

This optimized method successfully resolves amino acids, organic acids (TCA cycle), free sugars, phosphorylated intermediates, nucleotide mono-/di-/triphosphates (AMP, ADP, ATP), nucleotide sugars (ADP-Glc, UDP-Glc), and Acetyl-CoA in a single run. The method leverages both +ESI (predominantly for amino acids and Acetyl-CoA) and −ESI (for organic acids, sugars, and phosphorylated metabolites), proving that your LC mobile phase conditions accommodate both ion modes.

### 3.2. MS/MS fragmentation

Although Citric acid and Isocitric acid share an identical precursor ion (*m/z* 191) under negative ESI LC-MS/MS, considerable inconsistency exists across published literature regarding the selection of optimal product ions for quantification [30]. While the major fragment at m/z 111.0) is common to both isomers, various studies report disparate diagnostic transitions, such as *m/z* 155 and 73 for Isocitrate, versus *m/z* 87, 85, and 67 for Citric acid [31].

To resolve these literature discrepancies and rigorously evaluate each transition, we matched the published fragments with our experimental optimal MRM transitions generated via Guided Optimization MRM Infusion. Subsequently, to assess the accuracy and selectivity of each candidate fragment, calibration curves were established for both individual calibrants and mixed calibrant solutions of citric and isocitric acids. The percentage recovery of each acid in the individual solutions was calculated against its respective calibration curve of mixture (Table 1).

**Table 1.**
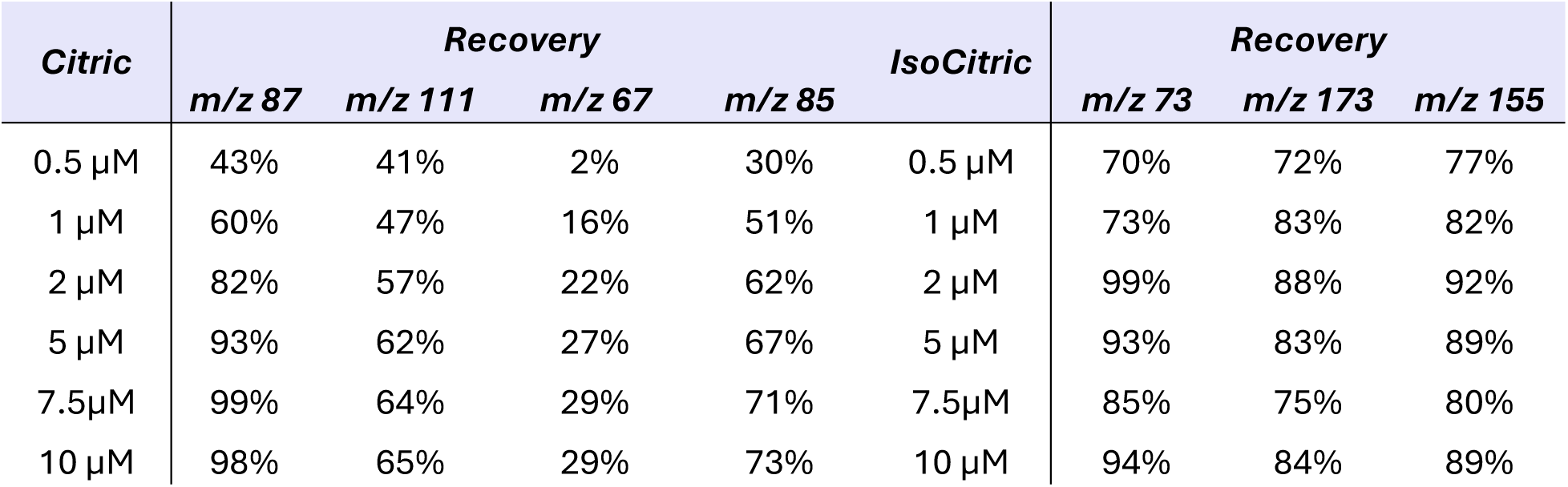
Recovery of single solution citric/isocitric versus mixture on different fragments.

| **Citric** | **Recovery** |  |  |  | **IsoCitric** | **Recovery** |  |  |
| --- | --- | --- | --- | --- | --- | --- | --- | --- |
|  | **m/z 87** | **m/z 111** | **m/z 67** | **m/z 85** |  | **m/z 73** | **m/z 173** | **m/z 155** |
| 0.5 μM | 43% | 41% | 2% | 30% | 0.5 μM | 70% | 72% | 77% |
| 1 μM | 60% | 47% | 16% | 51% | 1 μM | 73% | 83% | 82% |
| 2 μM | 82% | 57% | 22% | 62% | 2 μM | 99% | 88% | 92% |
| 5 μM | 93% | 62% | 27% | 67% | 5 μM | 93% | 83% | 89% |
| 7.5 μM | 99% | 64% | 29% | 71% | 7.5 μM | 85% | 75% | 80% |
| 10 μM | 98% | 65% | 29% | 73% | 10 μM | 94% | 84% | 89% |

For Citric acid: The m/z 87 fragment proved to be the best with high quantitative accuracy (82 – 99% recovery) within the 2 µM to 10 µM range. In contrast, alternative literature fragments failed to yield acceptable accuracy, m/z 67 suffered severe response attenuation amongst all. For Isocitric acid: The m/z 73 transition similarly provided the highest accuracy, achieving 85 – 99% recovery between 2 µM and 10 µM (peaking at 99% at 2 µM). Other candidate transitions, including m/z 155 (80 – 92%) and m/z 173 (75 – 88%), consistently underestimated the analyte concentration.

In plant and algal extracts, compounds like citric acid are present at millimolar levels, which far exceed the narrow linear range of standard calibration curves. This large concentration difference creates a severe risk of column saturation and injector carryover. Even with multi-solvent needle washes, physical carryover cannot be eliminated. Therefore, appropriate dilution factors must be applied to quantify abundant metabolites with carryover-free column and guard column.

### 3.3. Ion source temperature optimization

Optimizing the ion source temperature revealed a distinction between the free hexose sugars and the rest of metabolites included in the method. For the majority, represented by the amino acids and phosphorylated compounds (Fig. 1), higher temperatures significantly increased the signal intensity, reaching a maximum at 400 - 450 °C due to better desolvation. In contrast, the free hexoses (fructose and glucose) proved to be uniquely heat-sensitive, achieving their best signal at just 200 - 250 °C and thermal degradation starts when temperature was above 300 °C. Although an intermediate source temperature of 300 °C to 350 °C can be chosen as a compromise to maintain acceptable sensitivity for all analytes, in this work, metabolites were measured at two ion source temperatures as described in section 2.4.

**Figure 1.**
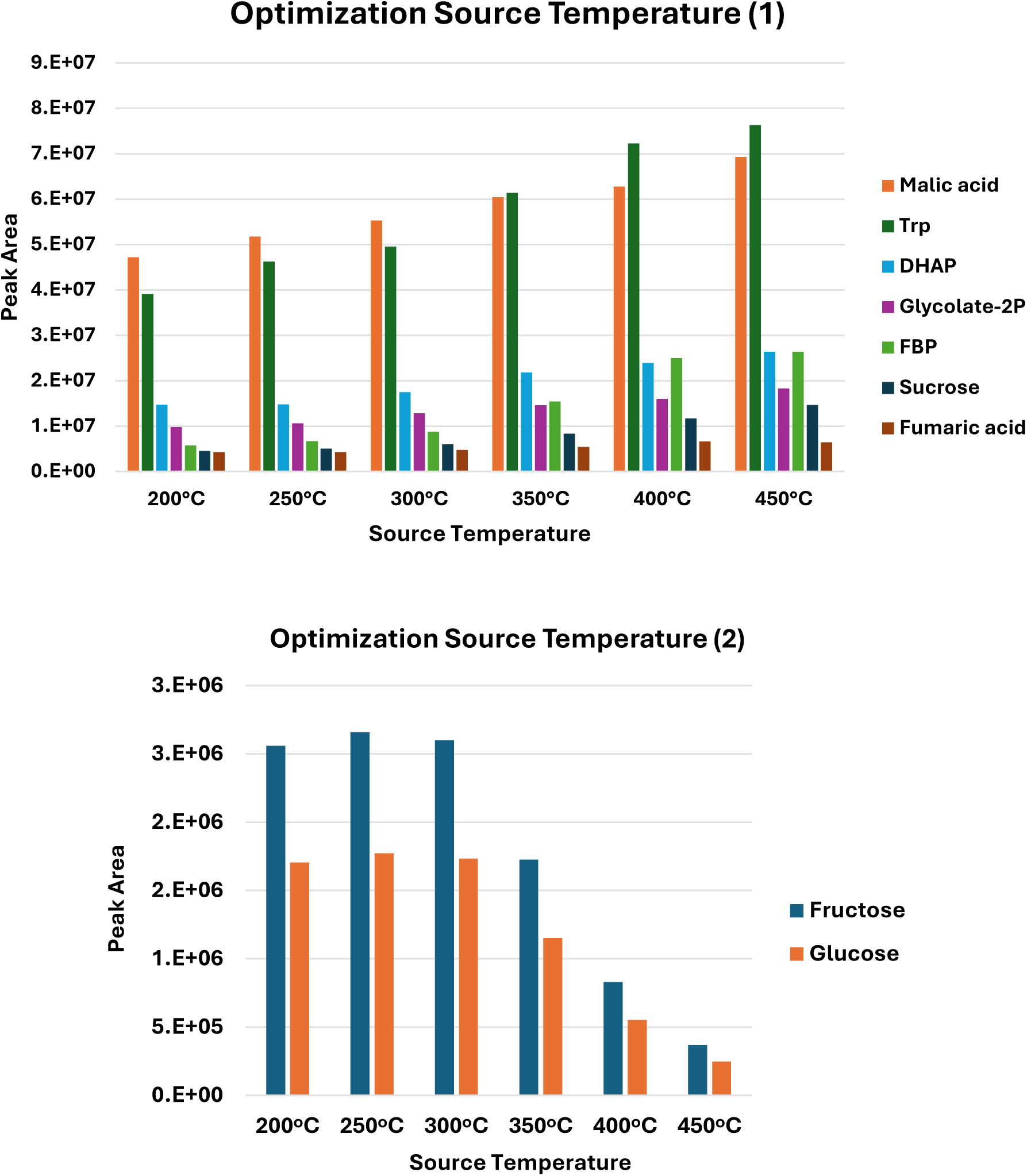
Ion source temperature optimization from 200 – 450°C (1) Peak area of representative amino acid, carboxylic acids and phosphorylated metabolites. (2) Peak area of fructose and glucose

### 3.4. Method validation

Method validation established high linearity (*R*^2^ ≥ 0.990) across a panel of more than 50 central metabolites spanning amino acids, organic acids, soluble sugars, Acetyl-CoA, and phosphorylated intermediates ranges spanning up to three orders of magnitude (Table 2). Operating with rapid polarity switching (+ESI/-ESI), the method demonstrated good retention time repeatability thanks to fluoropolymer mobile phase bottles and optimal concentration of modifier MDP. Over 40 runs, the relative standard deviations (RSD) remain below 1.3% for nearly all compounds, except glucose and histidine due to suboptimal chromatographic condition discussed in section 3.1.

**Table 2.** Overview of Retention time (RT) intraday repeatability, linearity ranges and limit of detection (LOQ) of metabolites.

| Compound | RT (min) | RSD (%) | Polarity | Min-LOQ (μM) | Max (μM) | *R2 | Compound | RT (min) | RSD (%) | Polarity | Min-LOQ (μM) | Max (μM) | *R2 |
| --- | --- | --- | --- | --- | --- | --- | --- | --- | --- | --- | --- | --- | --- |
| Trp | 1.7 | 0.5% | +ESI | 0.02 | 20 | 0.997 | 2-Ketoglutaric | 5.6 | 0.3% | -ESI | 1.00 | 15 | 0.999 |
| Phe | 1.8 | 0.5% | +ESI | 0.02 | 20 | 0.997 | Trehalose | 5.7 | 0.1% | -ESI | 0.20 | 15 | 0.996 |
| ILe | 1.9 | 0.4% | +ESI | 0.05 | 20 | 0.996 | Succinic | 5.8 | 0.3% | -ESI | 0.20 | 20 | 0.998 |
| Leu | 2.1 | 0.4% | +ESI | 0.05 | 15 | 0.997 | Fumaric | 6.3 | 0.2% | -ESI | 0.50 | 20 | 0.999 |
| Met | 2.3 | 0.4% | +ESI | 0.20 | 20 | 0.998 | Malic | 7.4 | 0.2% | -ESI | 0.10 | 20 | 0.994 |
| Val | 2.6 | 0.4% | +ESI | 0.50 | 20 | 0.996 | AMP | 8.0 | 0.2% | -ESI | 0.50 | 20 | 0.995 |
| Tyr | 2.6 | 0.3% | +ESI | 0.20 | 20 | 0.995 | DHAP | 8.1 | 0.6% | -ESI | 0.10 | 20 | 0.995 |
| Pro | 2.8 | 0.4% | +ESI | 0.02 | 20 | 0.997 | Glycerol-3P | 8.1 | 0.2% | -ESI | 1.00 | 15 | 0.995 |
| Ala | 3.8 | 0.2% | +ESI | 0.20 | 20 | 0.996 | Xylulose5P | 8.6 | 0.1% | -ESI | 0.50 | 20 | 0.991 |
| Thr | 4.1 | 0.3% | +ESI | 0.10 | 20 | 0.996 | Ribose5P | 9.0 | 0.1% | -ESI | 0.20 | 15 | 0.986 |
| Gly | 4.3 | 0.2% | +ESI | 2.00 | 20 | 0.993 | GAL3P | 9.3 | 0.6% | -ESI | 2.00 | 20 | 0.995 |
| Gln | 4.8 | 0.2% | +ESI | 0.20 | 20 | 0.997 | E4P | 9.4 | 0.5% | -ESI | 0.10 | 15 | 0.996 |
| Ser | 4.9 | 0.2% | +ESI | 0.50 | 20 | 0.996 | ADP-Glc | 9.4 | 0.3% | -ESI | 0.02 | 15 | 0.962 |
| Asn | 5.0 | 0.2% | +ESI | 0.20 | 15 | 0.996 | UDP-Glc | 10.1 | 0.3% | -ESI | 0.05 | 15 | 0.979 |
| Glu | 6.8 | 0.8% | +ESI | 2.00 | 20 | 0.997 | Sedo 7P | 10.3 | 0.1% | -ESI | 0.10 | 20 | 0.992 |
| His | 7.0 | 8.1% | +ESI | 0.50 | 15 | 0.997 | Glycolate-2P | 10.3 | 0.1% | -ESI | 1.00 | 20 | 0.997 |
| Asp | 7.1 | 0.2% | +ESI | 0.10 | 20 | 0.995 | PEP | 10.5 | 0.3% | -ESI | 0.01 | 20 | 0.999 |
| Acetyl-CoA | 9.4 | 0.2% | +ESI | 0.01 | 20 | 0.997 | 2PGA | 10.3 | 0.2% | -ESI | 0.10 | 15 | 0.991 |
| Arg | 9.2 | 0.5% | +ESI | 0.10 | 20 | 0.999 | 3PGA | 10.6 | 0.3% | -ESI | 0.50 | 20 | 0.999 |
| Orn | 9.7 | 0.1% | +ESI | 0.50 | 15 | 0.996 | ADP | 10.5 | 0.3% | -ESI | 0.10 | 20 | 0.998 |
| Lys | 9.9 | 0.2% | +ESI | 1.00 | 15 | 0.984 | IsoCitric | 10.7 | 0.2% | -ESI | 0.05 | 20 | 0.999 |
| Pyruvic | 1.2 | 0.8% | -ESI | 0.50 | 20 | 0.996 | Citric | 10.6 | 0.3% | -ESI | 0.50 | 10 | 0.982 |
| Glycolic | 2.3 | 0.5% | -ESI | 0.50 | 20 | 0.996 | ATP | 12.4 | 0.7% | -ESI | 0.01 | 15 | 0.998 |
| Fructose | 2.3 | 0.6% | -ESI | 0.50 | 20 | 0.997 | 6-PGluconic Acid | 12.6 | 0.2% | -ESI | 0.05 | 15 | 0.995 |
| Glucose | 2.8 | 1.7% | -ESI | 1.00 | 20 | 0.997 | RuBP | 13.7 | 0.1% | -ESI | 0.05 | 15 | 0.996 |
| Glyoxylic acid | 3.0 | 0.8% | -ESI | 0.50 | 15 | 0.996 | FBP | 13.9 | 0.1% | -ESI | 0.02 | 15 | 0.995 |
| Glyceric | 2.9 | 1.2% | -ESI | 0.10 | 20 | 0.998 | F6P | 9.7 | 0.1% | -ESI | 0.20 | 20 | 0.990 |
| Ascorbic Acid | 3.3 | 1.3% | -ESI | 2.00 | 20 | 0.968 | G1P | 10.1 | 0.1% | -ESI | 0.20 | 15 | 0.980 |
| Sucrose | 4.5 | 0.2% | -ESI | 0.20 | 20 | 0.993 | M6P | 10.3 | 0.6% | -ESI | 0.05 | 15 | 0.992 |
| Shikimate | 4.8 | 1.1% | -ESI | 1.00 | 15 | 0.996 | G6P | 10.9 | 0.7% | -ESI | 0.02 | 15 | 0.998 |

Regardless of the ion source temperature, unphosphorylated hexoses showed mutarotation issues. For glucose, slow mutarotation caused the peak to split into two separate peaks on both the ZIC-c and BEH Amide columns [15]. This peak doubling produced an unstable plateau rather than a distinct chromatographic apex, making it difficult to determine the exact RT and reducing RT repeatability (Figure S3).

The limit of Quantitation (LOQ) for each metabolite was defined as the lowest experimental concentration on the calibration curve demonstrating signal/noise (S/N) approx.10. Notably, multiple critical phosphorylated and metal-sensitive metabolites such as ATP, PEP, Ribulose-1,5-BP, and FBP achieved very good LOQ levels between 10 and 50 nM in negative mode, confirming that the passivation strategy successfully eliminated secondary column interactions without compromising ionization efficiency. Only a small subset shows marginally lower linearity (e.g., ADP-Glc, UDP-Glc, Ascorbic Acid, Citric acid and G1P), but they remain well within acceptable semi-quantitative bioanalytical limits.

### 3.5. Applications

#### Comparison of filters for *Chlamydomonas reinhardtii* harvest

To evaluate the impact of the harvesting filter materials, *Chlamydomonas reinhardtii* cultures were filtered using either glass fiber (GF/C) or nitrocellulose (NC) membranes. The content (nmol g^−1^ Dry weight (DW)) of metabolites is shown in Table 3. Both filter materials show exceptional agreement (RSD ≤ 3%) for abundant and stable amino acids such as Thr, Leu, Ser, Glu and carboxylic acids including malic, pyruvic as well phosphorylated intermediates such as G6P and FBP. Meanwhile, several metabolites, including Ascorbic acid, Glyoxylic acid, Shikimic acid, Fructose, Acetyl-CoA, E4P, and GAL3P, were not detected in either filter type, while glycolate 2P was detected but could not be quantified.

**Table 3:** Content of metabolites (nmol g^−1^ Dry weight (DW)) found in Chlamydomonas reinhardtii (n = 3) using glass fiber (GF/C) or nitrocellulose (NC) filter.

| Compound | Category | GF/C | NC | RSD (%) |
| --- | --- | --- | --- | --- |
| Thr | Amino acid | 518.8 ± 14.0 | 513.6 ± 33.2 | 1% |
| Leu | Amino acid | 212.1 ± 18.3 | 208.9 ± 15.1 | 1% |
| Ser | Amino acid | 1770.8 ± 207.7 | 1726.4 ± 85.0 | 2% |
| Glu | Amino acid | 33920.4 ± 897.6 | 35363.7 ± 1970.6 | 3% |
| Asn | Amino acid | 436.3 ± 7.9 | 459.4 ± 19.3 | 4% |
| His | Amino acid | 365.7 ± 54.0 | 344.6 ± 69.1 | 4% |
| Arg | Amino acid | 796.7 ± 11.3 | 749.6 ± 87.9 | 4% |
| Tyr | Amino acid | 233.6 ± 12.3 | 251.0 ± 24.0 | 5% |
| Gly | Amino acid | 623.6 ± 49.9 | 678.9 ± 54.7 | 6% |
| Lys | Amino acid | 627.7 ± 55.9 | 571.9 ± 28.5 | 7% |
| Val | Amino acid | 434.0 ± 11.2 | 508.4 ± 26.0 | 11% |
| Trp | Amino acid | 74.7 ± 4.3 | 88.4 ± 4.7 | 12% |
| Pro | Amino acid | 235.7 ± 6.1 | 197.5 ± 3.8 | 12% |
| Ala | Amino acid | 2505.3 ± 60.3 | 1970.7 ± 224.7 | 17% |
| Phe | Amino acid | 247.4 ± 9.8 | 330.2 ± 21.4 | 20% |
| Orn | Amino acid | 248.2 ± 28.2 | 182.8 ± 21.8 | 21% |
| Gln | Amino acid | 1952.4 ± 81.7 | 2715.5 ± 365.5 | 23% |
| Met | Amino acid | 181.1 ± 21.8 | 129.2 ± 43.4 | 24% |
| Ile | Amino acid | 320.9 ± 119.1 | 465.9 ± 31.9 | 26% |
| Asp | Amino acid | 1181.5 ± 57.3 | 593.9 ± 60.5 | 47% |
| Ascorbic Acid | Carboxylic acid | ND | ND | NA |
| Glyoxylic acid | Carboxylic acid | ND | ND | NA |
| Shikimic acid | Carboxylic acid | ND | ND | NA |
| Malic acid | Carboxylic acid | 3218.1 ± 355.6 | 3242.6 ± 133.8 | 1% |
| Pyruvic acid | Carboxylic acid | 1531.1 ± 209.2 | 1470.4 ± 455.5 | 3% |
| Isocitric acid | Carboxylic acid | 56.9 ± 4.4 | 61.1 ± 9.2 | 5% |
| Glyceric acid | Carboxylic acid | 146.6 ± 45.6 | 123.9 ± 3.1 | 12% |
| Fumaric acid | Carboxylic acid | 396.2 ± 58.4 | 330.0 ± 24.5 | 13% |
| Succinic acid | Carboxylic acid | 1772.5 ± 274.9 | 1412.2 ± 55.4 | 16% |
| Citric acid | Carboxylic acid | 2328.3 ± 87.0 | 1641.9 ± 152.7 | 24% |
| 6-PGluconic Acid | Carboxylic acid | 341.1 ± 70.5 | 217.0 ± 51.5 | 31% |
| 2-Ketoglutaric acid | Carboxylic acid | 183.4 ± 24.8 | 306.3 ± 70.0 | 36% |
| Glycolic acid | Carboxylic acid | 4543.4 ± 1613.7 | 1486.0 ± 348.1 | 72% |
| AMP | Nucleotide / Phosphate | 2632.7 ± 721.6 | 3090.3 ± 60.3 | 11% |
| ATP | Nucleotide / Phosphate | 640.1 ± 143.6 | 1420.9 ± 258.2 | 54% |
| ADP | Nucleotide / Phosphate | 1068.5 ± 143.7 | 2736.1 ± 527.8 | 62% |
| ADP-Glc | Nucleotide Sugar | 8.7 ± 0.5 | 15.2 ± 0.9 | 39% |
| UDP-Glc | Nucleotide Sugar | 373.0 ± 13.5 | 737.3 ± 406.7 | 46% |
| **E4P** | Phosphorylated Metabolite | ND | ND | NA |
| **GAL3P** | Phosphorylated Metabolite | ND | ND | NA |
| **Glycolate-2P** | Phosphorylated Metabolite | NQ | NQ | NA |
| **FBP** | Phosphorylated Metabolite | 2058.9 ± 641.0 | 2052.1 ± 320.2 | 0% |
| **G6P** | Phosphorylated Metabolite | 1095.2 ± 125.9 | 1082.1 ± 92.3 | 1% |
| **3PGA** | Phosphorylated Metabolite | 6451.2 ± 441.5 | 7068.8 ± 231.8 | 6% |
| **Ribose5P** | Phosphorylated Metabolite | 249.1 ± 33.1 | 204.2 ± 29.0 | 14% |
| **Xy5P+Ru5P<sup>1</sup>** | Phosphorylated Metabolite | 919.9 ± 160.7 | 753.9 ± 113.4 | 14% |
| **DHAP** | Phosphorylated Metabolite | 1134.0 ± 67.2 | 854.0 ± 94.6 | 20% |
| **PEP** | Phosphorylated Metabolite | 554.9 ± 159.7 | 746.2 ± 216.1 | 21% |
| **RuBP** | Phosphorylated Metabolite | 621.1 ± 138.8 | 843.2 ± 78.9 | 21% |
| **M6P** | Phosphorylated Metabolite | 1268.1 ± 67.9 | 929.1 ± 182.9 | 22% |
| **G1P** | Phosphorylated Metabolite | 520.8 ± 74.9 | 718.2 ± 230.9 | 23% |
| **F6P** | Phosphorylated Metabolite | 1614.6 ± 661.8 | 1074.9 ± 64.6 | 28% |
| **Glycerol-3P** | Phosphorylated Metabolite | 1907.4 ± 193.6 | 1106.4 ± 167.0 | 38% |
| **S7P** | Phosphorylated Metabolite | 448.1 ± 88.5 | 822.7 ± 103.6 | 42% |
| **2PGA** | Phosphorylated Metabolite | 235.2 ± 134.3 | 711.4 ± 332.5 | 71% |
| **Fructose** | Sugar | ND | ND | NA |
| **Glucose** | Sugar | 35924.6 ± 3993.4 | 65819.2 ± 11542.2 | 42% |
| **Sucrose** | Sugar | 14.4 ± 11.8 | 29.0 ± 10.1 | 48% |
| **Trehalose** | Sugar | 86.7 ± 9.0 | 70.7 ± 10.6 | 14% |
| **Acetyl-CoA** | Coenzyme | ND | ND | NA |

ND: Not detected, ND: not quantifiable, NA: Not applicable
<sup>1</sup> Xy5P+Ru5P were quantified together based on Xy5P standard.

For metabolites with RSD > 15% between the two filters, GF/C gave much higher results for four amino acids (Ala, Orn, Met, and especially Asp, which was twice as high as on NC), as well as five out of six carboxylic acids, except for 2-ketoglutaric acid. In addition, the levels of all adenine nucleotides and nucleotide sugars showed a strong shift on NC filter with RSDs of 39 – 62% alongside key glycolytic and pentose phosphate pathway intermediates such as 2PGA, S7P and glycerol-3-phosphate with RSDs of 38 – 71%.

Rather than genuine biological variation, the observed deviations on nitrocellulose (NC) membranes were largely driven by chemical instability in organic extraction solvents (chloroform/methanol). This degradation introduced substantial background noise, ion suppression, and ghost peaks across several targeted MRM transitions. Although NC filters were previously used to collect freshwater cyanobacteria [32]; those extracts underwent chemical derivatization prior to analysis, which likely minimized matrix effects and background interference.

In contrast, borosilicate GF/C filters provided complete solvent compatibility, absence of leachable polymer artifacts, and reproducible peak baselines, establishing GF/C as the preferred filter matrix for reliable metabolomics profiling in *Chlamydomonas reinhardtii*.

#### Preliminary extraction screening for *Arabidopsis thaliana*

During initial development, four extraction conditions were evaluated using 50 mg of homogenized *Arabidopsis thaliana*: liquid–liquid extraction (LLE) adapted from the algal protocol using three freeze–thaw cycles and collected to lyophilize in either 1.5 mL tubes (condition 1) or 5 mL tubes (condition 2), LLE with incubation at −20°C for 2 h instead of freeze–thawing (condition 3), and a solid–liquid extraction (SLE) method using 225 µL of Milli-Q/Acetonitrile/Isopropanol (2:3:3, v/v/v) [8] agitated at 4°C for 30 min (condition 4), with each condition spiked with 25 µL of 100 µM Ribitol.

Using 50 mg of starting material introduced substantial analytical drawbacks for LC-MS/MS analysis. The high matrix load caused LC column clogging and severe ion suppression on RuBP, necessitating a reduction in initial sample biomass to 20 mg as optimal. Incubation at −20°C for 2 h (condition 3) slowed down the extraction and yielded lower signal intensities than the freeze thaw approach. Because conditions 1 and 2 showed no significant difference, condition 1 was chosen to save time and reduce labor by avoiding the 2 h delay and using standard 1.5 mL tubes.

Among the tested conditions, the SLE in condition 4 gave the poorest chromatographic with distorted peaks and very weak signals results (Figure S4). Due to lack of nonpolar phase in condition 4 to separate lipids, extracts retained a strong green color from co-extracted chlorophyll pigments. While this solvent mixture works well for GC-MS workflows with derivatization, it is unsuitable for direct HILIC-MS/MS because it caused severe matrix interferences and suppressed signals for nearly all phosphorylated metabolites, including ATP, AMP, glycerol 3-phosphate, 3PGA, S7P, FBP and sugar mono phosphates.

#### Effect of pestle (for cell strainer) and cooling Arabidopsis extraction

Because grinding sample with a liquid-nitrogen-cooled porcelain mortar and pestle already completely disrupted the cells during harvesting, additional freeze–thaw cycles were unnecessary. To determine whether further manual homogenization and prolonged low-temperature incubation were needed, four conditions were compared: with or without micro pestle grinding, and with or without an extended cooling period (Figure 2). Overall, the total quantified content across six metabolite groups did not exhibit significant differences among the treatments (Table S4).

**Figure 2.**
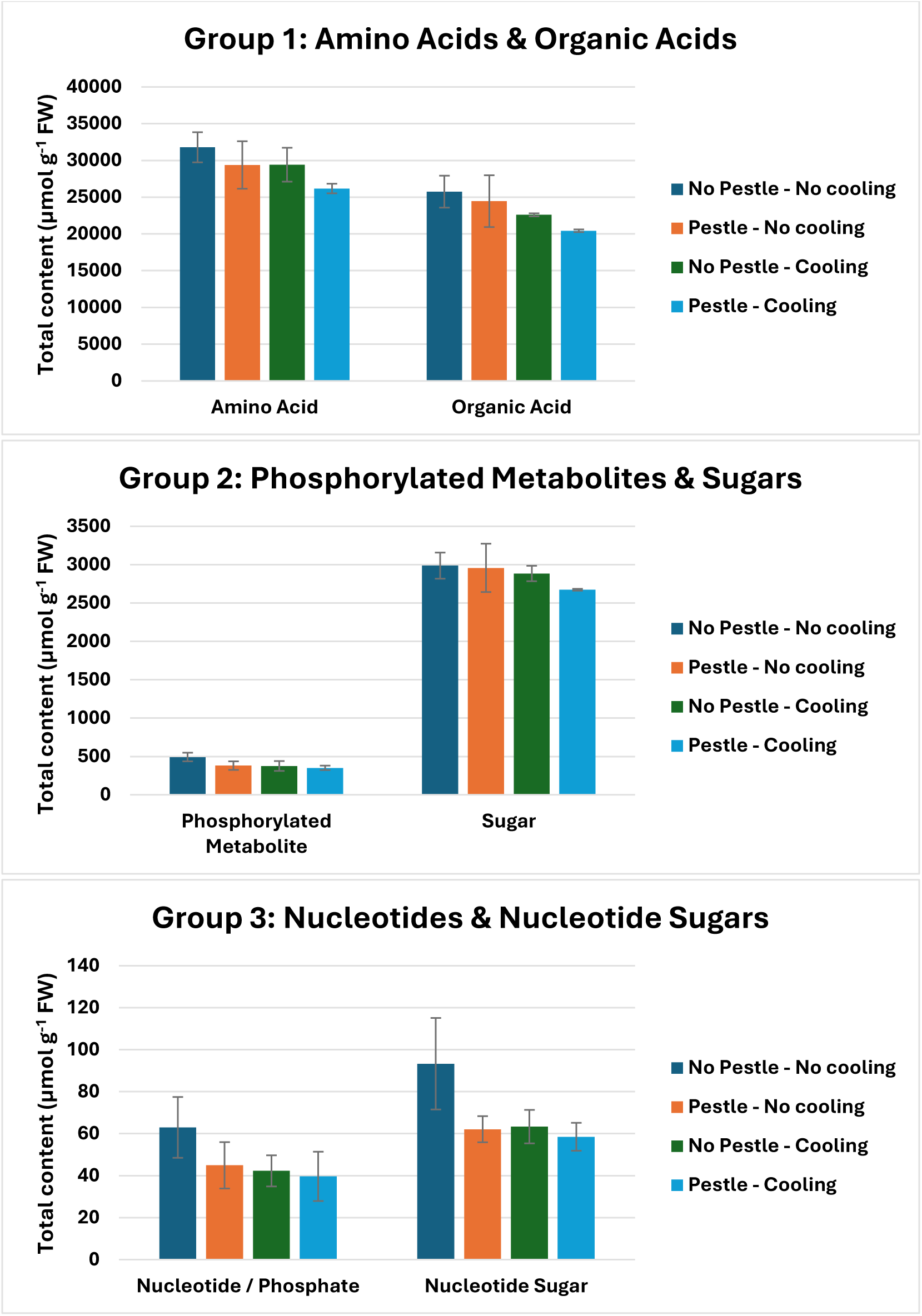
Content (µmol g^−1^ Fresh Weight (FW)) of 3 metabolite groups in Arabidopsis thaliana under effect of micro pestle and extended cooling (n=3)

Across all groups, the “No Pestle, No Cooling” condition consistently yielded the highest metabolite contents. Additional pestle grinding and extended cooling did not improve extraction efficiency, only adding hands on time. Therefore, both steps were omitted.

Overall, in *Arabidopsis*, the most abundant metabolites were glutamate among amino acids, fumaric and citric acids among organic acids, and glucose and sucrose among sugars. In contrast, phosphorylated compounds and energy molecules were detected at much lower physiological levels, including ATP, ADP, AMP, and key pathway intermediates like F6P, DHAP, and 3PGA. Only Acetyl-CoA, E4P, and glycolate-2P were not detected.

## 4. Conclusions

In this work, we established a robust, sensitive, and reproducible HILIC-MS/MS workflow for the quantitative profiling of central carbon metabolism in photosynthetic organisms. By coupling a bioinert BEH Amide stationary phase with an alkaline mobile phase pH 9.0 containing a low salt concentration (10 mM ammonium acetate) and 75 µM medronic acid in mobile phase A, we successfully resolved metal-chelation bottlenecks for polyphosphorylated sugar phosphates and nucleotides without subjecting the LC-MS hardware to heavy salt deposition or high column temperature. The method achieved baseline separation for major positional isomers and provided excellent peak symmetry by collapsing sugar phosphate anomers. Successfully applied to quantify nearly 60 central intermediates in *Chlamydomonas reinhardtii* and *Arabidopsis thaliana*, this analytical framework provides a reliable foundation for downstream metabolic flux analysis and functional plant metabolomics.

## Acknowledgement

This project was supported by the European Research Council (ERC) under the European Union’s Horizon Europe research and innovation programme (grant agreement No. 101076560, project FRIDOM), and the German Research Foundation (DFG, Grant INST 248/399-1 FUGG). OG was supported by the German Research Foundation (DFG, project no. SCHR 617/13-1 to HT). GO was supported by the German Research Foundation (DFG, TRR175/A09). NKNB would like to thank PD Dr. Franziska Fichtner (Heinrich Heine University Düsseldorf) for kindly providing the reference standard.

## References

[1] Treves, H. et al. (2021). Carbon flux through photosynthesis and central carbon metabolism show distinct patterns between algae, C3 and C4 plants. Nature Plants. 10.1038/s41477-021-01042-5.

[2] Cocuron, J.-C. et al. (2014). Targeted Metabolomics of Physaria fendleri, an Industrial Crop Producing Hydroxy Fatty Acids. Plant and Cell Physiology. 10.1093/pcp/pcu011.

[3] Xu, Y. and Fu, X. (2022). Reprogramming of Plant Central Metabolism in Response to Abiotic Stresses: A Metabolomics View. International Journal of Molecular Sciences. 10.3390/ijms23105716.

[4] Wang, W. et al. (2022). Sucrose synthase activity is not required for cellulose biosynthesis in Arabidopsis. The Plant Journal. 10.1111/tpj.15752.

[5] Timm, S. and Arrivault, S. (2021). Regulation of Central Carbon and Amino Acid Metabolism in Plants. Plants. 10.3390/plants10030430.

[6] Yan, S. et al. (2022). Recent advances in proteomics and metabolomics in plants. Molecular Horticulture. 10.1186/s43897-022-00038-9.

[7] Gullberg, J. et al. (2004). Design of experiments: an efficient strategy to identify factors influencing extraction and derivatization of Arabidopsis thaliana samples in metabolomic studies with gas chromatography/mass spectrometry. Analytical Biochemistry. 10.1016/j.ab.2004.04.037.

[8] Durand, T.C. et al. (2019). Combined Proteomic and Metabolomic Profiling of the Arabidopsis thaliana vps29 Mutant Reveals Pleiotropic Functions of the Retromer in Seed Development. International Journal of Molecular Sciences. 10.3390/ijms20020362.

[9] Tang, D. et al. (2016). HILIC-MS for metabolomics: An attractive and complementary approach to RPLC-MS. Mass Spectrometry Reviews. 10.1002/mas.21445.

[10] Arrivault, S. et al. (2009). Use of reverse-phase liquid chromatography, linked to tandem mass spectrometry, to profile the Calvin cycle and other metabolic intermediates in Arabidopsis rosettes at different carbon dioxide concentrations. The Plant Journal. 10.1111/j.1365-313X.2009.03902.x.

[11] Koley, S. et al. (2022). An efficient LC-MS method for isomer separation and detection of sugars, phosphorylated sugars, and organic acids. Journal of Experimental Botany. 10.1093/jxb/erac062.

[12] Stafsnes, M.H. et al. (2018). Improved phosphometabolome profiling applying isotope dilution strategy and capillary ion chromatography-tandem mass spectrometry. Journal of Chromatography B. 10.1016/j.jchromb.2018.02.004.

[13] Serafimov, K. and Lämmerhofer, M. (2024). Comprehensive Coverage of Glycolysis and Pentose Phosphate Metabolic Pathways by Isomer-Selective Accurate Targeted Hydrophilic Interaction Liquid Chromatography-Tandem Mass Spectrometry Assay. Analytical Chemistry. 10.1021/acs.analchem.4c03490.

[14] Xu, K. et al. (2025). Development and Validation of Targeted Metabolomics Methods Using Liquid Chromatography–Tandem Mass Spectrometry (LC-MS/MS) for the Quantification of 235 Plasma Metabolites. Molecules. 10.3390/molecules30030706.

[15] Hosseinkhani, F. et al. (2022). Systematic Evaluation of HILIC Stationary Phases for Global Metabolomics of Human Plasma. Metabolites. 10.3390/metabo12020165.

[16] Langová, A. et al. (2026). Enhancing metabolite coverage using dedicated mobile phases for individual polarity modes in HILIC-MS. Analytical and Bioanalytical Chemistry. 10.1007/s00216-025-06189-0.

[17] Serafimov, K. et al. (2025). UCL-MetIsoLib: A Public High-Resolution Tandem Mass Spectrometry Library for HILIC-Based Isomer-Resolved Profiling of Glycolysis, Central Carbon Metabolism, and Beyond in Urine, Plasma, Tissues, Cells, and Patient-Derived Organoids. Analytical Chemistry. 10.1021/acs.analchem.5c03390.

[18] McCalley, D.V. (2022). Influence of metals in the column or instrument on performance in hydrophilic interaction liquid chromatography. Journal of Chromatography A. 10.1016/j.chroma.2021.462751.

[19] Shen, D. et al. (2026). Integrated HILIC-MS/MS strategy for targeted analysis of key central carbon metabolism metabolites: Resolving isomers and overcoming metal chelation in complex biological matrices. Talanta. 10.1016/j.talanta.2026.129605.

[20] Hsiao, J.J. et al. (2018). Improved LC/MS Methods for the Analysis of Metal-Sensitive Analytes Using Medronic Acid as a Mobile Phase Additive. Analytical Chemistry. 10.1021/acs.analchem.8b02100.

[21] Hsu, J. et al. (2023). Carnitine octanoyltransferase is important for the assimilation of exogenous acetyl-L-carnitine into acetyl-CoA in mammalian cells. Journal of Biological Chemistry. 10.1016/j.jbc.2022.102848.

[22] Schwaiger-Haber, M. et al. (2023). Using mass spectrometry imaging to map fluxes quantitatively in the tumor ecosystem. Nature Communications. 10.1038/s41467-023-38403-x.

[23] Chan, J.K. et al. (2023). Protocol for mapping the metabolome and lipidome of medulloblastoma cells using liquid chromatography-mass spectrometry. STAR Protocols. 10.1016/j.xpro.2023.102736.

[24] Keller, P. et al. (2022). Generation of an Escherichia coli strain growing on methanol via the ribulose monophosphate cycle. Nature Communications. 10.1038/s41467-022-32744-9.

[25] Gurler, S.B. et al. (2023). HER2 overexpression initiates breast tumorigenesis non-cell-autonomously by inducing oxidative stress in the tissue microenvironment. 10.1101/2023.08.25.554770.

[26] Scholefield, M. et al. (2023). Multi-regional alterations in glucose and purine metabolic pathways in the Parkinson’s disease dementia brain. npj Parkinson’s Disease. 10.1038/s41531-023-00488-y.

[27] Geffen, O. et al. (2023). ^13^CO_2_-labelling and Sampling in Algae for Flux Analysis of Photosynthetic and Central Carbon Metabolism. BIO-PROTOCOL. 10.21769/BioProtoc.4808.

28. Khold-Atila, A. (2024). Exploring C. Ohadii’s metabolic flexibility - studying the physiological state function and the metabolic pathways along different growth phases.

[29] Serafimov, K. et al. (2024). Solving the retention time repeatability problem of hydrophilic interaction liquid chromatography. Journal of Chromatography A. 10.1016/j.chroma.2024.465060.

30. Al Kadhi, O., et al. (2017). Development of a LC-MS/MS Method for the Simultaneous Detection of Tricarboxylic Acid Cycle Intermediates in a Range of Biological Matrices. Journal of Analytical Methods in Chemistry. 10.1155/2017/5391832.

[31] Giacomello, G. et al. (2022). Isotopic tracing of glucose metabolites in human monocytes to assess changes in inflammatory conditions. STAR Protocols. 10.1016/j.xpro.2022.101715.

[32] Perin, G. et al. (2021). Calm on the surface, dynamic on the inside. Molecular homeostasis of *Anabaena* sp. PCC 7120 nitrogen metabolism. Plant, Cell & Environment. 10.1111/pce.14034.

