## Supplementary Information for "Targeted Quantitative Analysis of Polar and Phosphorylated Intermediates of Algal and Plant Primary Carbon Metabolism Using an Optimized HILIC-Amide Tandem MS Workflow"

*Table S1. List of metabolites and their abbreviations*

| No. | Full Name | Abbreviation | Category |
|---|---|---|---|
| 1 | Tryptophan | Trp | Amino Acid |
| 2 | Phenylalanine | Phe | Amino Acid |
| 3 | Isoleucine | ILe | Amino Acid |
| 4 | Leucine | Leu | Amino Acid |
| 5 | Methionine | Met | Amino Acid |
| 6 | Valine | Val | Amino Acid |
| 7 | Tyrosine | Tyr | Amino Acid |
| 8 | Proline | Pro | Amino Acid |
| 9 | Hydroxyproline | HPro | Amino Acid |
| 10 | Alanine | Ala | Amino Acid |
| 11 | Threonine | Thr | Amino Acid |
| 12 | Glycine | Gly | Amino Acid |
| 13 | Glutamine | Gln | Amino Acid |
| 14 | Serine | Ser | Amino Acid |
| 15 | Asparagine | Asn | Amino Acid |
| 16 | Glutamic acid | Glu | Amino Acid |
| 17 | Histidine | His | Amino Acid |
| 18 | Aspartic acid | Asp | Amino Acid |
| 19 | Arginine | Arg | Amino Acid |
| 20 | Ornithine | Orn | Amino Acid |
| 21 | Lysine | Lys | Amino Acid |

| No. | Full Name | Abbreviation | Category |
|---|---|---|---|
| 22 | Adenosine monophosphate | AMP | Nucleotide / Phosphate |
| 23 | Adenosine diphosphate | ADP | Nucleotide / Phosphate |
| 24 | Adenosine triphosphate | ATP | Nucleotide / Phosphate |
| 25 | ADP-glucose | ADP-Glc | Nucleotide Sugar |
| 26 | UDP-glucose | UDP-Glc | Nucleotide Sugar |
| 27 | Pyruvic acid | NA | Organic Acid |
| 28 | Glycolic acid | NA | Organic Acid |
| 29 | Glyceric acid | NA | Organic Acid |
| 30 | Shikimic acid | NA | Organic Acid |
| 31 | 2-Ketoglutaric acid | NA | Organic Acid |
| 32 | Succinic acid | NA | Organic Acid |
| 33 | Fumaric acid | NA | Organic Acid |
| 34 | Malic acid | NA | Organic Acid |
| 35 | Glyoxylic acid | NA | Organic Acid |
| 36 | Citric acid | NA | Organic Acid |
| 37 | Isocitric acid | NA | Organic Acid |
| 38 | Dihydroxyacetone phosphate | DHAP | Phosphorylated Metabolite |
| 39 | Glycerol 3-phosphate | Glycerol-3P | Phosphorylated Metabolite |
| 40 | Xylulose 5-phosphate | Xy5P | Phosphorylated Metabolite |
| 41 | Ribulose 5-phosphate | Ru5P | Phosphorylated Metabolite |
| 42 | Ribose 5-phosphate | R5P | Phosphorylated Metabolite |
| 43 | Glyceraldehyde 3-phosphate | GAL3P | Phosphorylated Metabolite |
| 44 | Sedoheptulose 7-phosphate | S7P | Phosphorylated Metabolite |
| 45 | 6-Phosphogluconic acid | 6-PGluconic Acid | Phosphorylated Metabolite |
| 46 | Ribulose 1,5-bisphosphate | RuBP | Phosphorylated Metabolite |
| 47 | Fructose 1,6-bisphosphate | FBP | Phosphorylated Metabolite |
| 48 | Fructose 6-phosphate | F6P | Phosphorylated Metabolite |
| 49 | Glucose 1-phosphate | G1P | Phosphorylated Metabolite |
| 50 | Mannose 6-phosphate | M6P | Phosphorylated Metabolite |
| 51 | Glucose 6-phosphate | G6P | Phosphorylated Metabolite |
| 52 | 2-Phosphoglycolate | Glycolate-2P | Phosphorylated Metabolite |
| 53 | Phosphoenolpyruvate | PEP | Phosphorylated Metabolite |
| 54 | 2-Phosphoglycerate | 2PGA | Phosphorylated Metabolite |
| 55 | 3-Phosphoglycerate | 3PGA | Phosphorylated Metabolite |
| 56 | Fructose | NA | Hexose |

| No. | Full Name | Abbreviation | Category |
|---|---|---|---|
| 57 | Glucose | NA | Hexose |
| 58 | Sucrose | NA | Sugar |
| 59 | Trehalose | NA | Sugar |

*NA: not applicable*

*Table S2. Positive mode transition table*

| Compound | Type | Retention time | Q1 mass | Q3 mass | DP | EP | CE | CXP |
|---|---|---|---|---|---|---|---|---|
| Trp | Quantifier | 1.7 | 205 | 188 | 5 | 10 | 16 | 11 |
| Trp | Qualifier | 1.7 | 205 | 146 | 5 | 10 | 25 | 12 |
| Phe | Quantifier | 1.8 | 166 | 120 | 55 | 10 | 26 | 15 |
| Phe | Qualifier | 1.8 | 166 | 131 | 55 | 10 | 26 | 15 |
| Ile | Quantifier | 1.9 | 132 | 86 | 25 | 10 | 14 | 11 |
| Ile | Qualifier | 1.9 | 132 | 43 | 31 | 10 | 25 | 10 |
| Leu | Quantifier | 2.1 | 132 | 86 | 25 | 10 | 14 | 11 |
| Leu | Qualifier | 2.1 | 132 | 69 | 31 | 10 | 25 | 10 |
| Met | Quantifier | 2.2 | 150 | 133 | 20 | 10 | 15 | 11 |
| Met | Qualifier | 2.2 | 150 | 104 | 46 | 10 | 15 | 12 |
| NVa | Quantifier | 2.3 | 118 | 72 | 30 | 10 | 17 | 7 |
| NVa | Qualifier | 2.3 | 118 | 30 | 30 | 10 | 26 | 7 |
| Val | Quantifier | 2.5 | 118 | 72 | 30 | 10 | 17 | 7 |
| Tyr | Quantifier | 2.6 | 182 | 165 | 10 | 10 | 14 | 19 |
| Tyr | Qualifier | 2.6 | 182 | 147 | 10 | 10 | 14 | 19 |
| Pro | Quantifier | 2.8 | 116 | 70 | 15 | 10 | 20 | 6 |
| Ala | Quantifier | 3.7 | 90 | 44 | 20 | 10 | 25 | 6 |
| Thr | Quantifier | 4.1 | 120 | 74 | 30 | 10 | 15 | 10 |
| Thr | Qualifier | 4.1 | 120 | 102 | 30 | 10 | 5 | 10 |
| Gly | Quantifier | 4.3 | 76 | 76 | 40 | 10 | 10 | 6 |
| Gly | Qualifier | 4.3 | 76 | 30 | 40 | 10 | 20 | 10 |
| Gln | Quantifier | 4.7 | 147 | 130 | 5 | 10 | 12 | 12 |
| Gln | Qualifier | 4.7 | 147 | 84 | 26 | 10 | 23 | 10 |
| Ser | Quantifier | 4.8 | 106 | 60 | 15 | 10 | 27 | 9 |
| Asn | Quantifier | 5.0 | 133 | 74 | 10 | 10 | 23 | 5 |
| Asn | Qualifier | 5.0 | 133 | 87 | 10 | 10 | 23 | 5 |
| Glu | Quantifier | 6.6 | 148 | 84 | 21 | 10 | 23 | 10 |
| Glu | Qualifier | 6.6 | 148 | 102 | 21 | 10 | 23 | 10 |
| His | Quantifier | 7.0 | 156 | 110 | 50 | 10 | 21 | 16 |
| Asp | Quantifier | 7.1 | 134 | 74 | 10 | 10 | 19 | 10 |
| Asp | Qualifier | 7.1 | 134 | 88 | 10 | 10 | 19 | 10 |
| Acetyl-CoA | Quantifier | 9.1 | 810 | 303 | 55 | 10 | 44 | 17 |
| Acetyl-CoA | Qualifier | 9.1 | 810 | 428 | 55 | 10 | 35 | 30 |
| Cystine | Quantifier | 9.2 | 241 | 152 | 30 | 10 | 20 | 14 |
| Cystine | Qualifier | 9.2 | 241 | 120 | 30 | 10 | 28 | 14 |
| Arg | Quantifier | 9.5 | 175 | 70 | 10 | 10 | 32 | 8 |
| Arg | Qualifier | 9.5 | 175 | 116 | 10 | 10 | 24 | 18 |
| Orn | Quantifier | 9.7 | 133 | 70 | 41 | 10 | 23 | 8 |
| Lys | Quantifier | 9.9 | 147 | 84 | 26 | 10 | 23 | 10 |

*Table S3. Negative mode transition table*

| Compound | Type | Retention time | Q1 mass | Q3 mass | DP | EP | CE | CXP |
|---|---|---|---|---|---|---|---|---|
| Pyruvic | Quantifier | 1.1 | 87 | 43 | -5 | -10 | -12 | -5 |
| Ribitol | Quantifier | 1.9 | 151 | 89 | -20 | -10 | -16 | -7 |
| Ribitol | Qualifier | 1.9 | 151 | 71 | -20 | -10 | -21 | -8 |
| Glycolic acid | Quantifier | 2.2 | 75 | 47 | -40 | -10 | -14 | -11 |
| Glycolic acid | Qualifier | 2.2 | 75 | 45 | -40 | -10 | -14 | -11 |
| Fructose | Quantifier | 2.5 | 179 | 89 | -40 | -10 | -12 | -11 |
| Fructose | Qualifier | 2.5 | 179 | 71 | -40 | -10 | -12 | -35 |
| Glucose | Quantifier | 2.8 | 179 | 89 | -40 | -10 | -12 | -11 |
| Glucose | Qualifier | 2.8 | 179 | 71 | -40 | -10 | -12 | -35 |
| Glyoxylic acid | Quantifier | 2.8 | 73 | 45 | -35 | -10 | -11 | -8 |
| Glyceric acid | Quantifier | 2.9 | 105 | 75 | -15 | -10 | -16 | -7 |
| Glyceric acid | Qualifier | 2.9 | 105 | 59 | -15 | -10 | -16 | -7 |
| Ascorbic acid | Quantifier | 3.2 | 175 | 87 | -15 | -10 | -26 | -10 |
| Ascorbic acid | Qualifier | 3.2 | 175 | 115 | -15 | -10 | -15 | -10 |
| Sucrose | Quantifier | 4.5 | 341 | 59 | -75 | -10 | -50 | -11 |
| Sucrose | Qualifier | 4.5 | 341 | 89 | -75 | -10 | -40 | -11 |
| Shikimic acid | Quantifier | 4.8 | 173 | 93 | -30 | -10 | -22 | -11 |
| Shikimic acid | Qualifier | 4.8 | 173 | 111 | -30 | -10 | -22 | -11 |
| 2-Ketoglutaric acid | Quantifier | 5.1 | 145 | 101 | -10 | -10 | -12 | -4 |
| 2-Ketoglutaric acid | Qualifier | 5.1 | 145 | 57 | -10 | -10 | -15 | -26 |
| Trehalose | Quantifier | 5.2 | 341 | 59 | -75 | -10 | -50 | -11 |
| Trehalose | Qualifier | 5.2 | 341 | 71 | -125 | -10 | -48 | -7 |
| Succinic acid | Quantifier | 5.7 | 117 | 73 | -15 | -10 | -17 | -10 |
| Succinic acid | Qualifier | 5.7 | 117 | 99 | -15 | -10 | -14 | -9 |
| Fumaric acid | Quantifier | 6.3 | 115 | 71 | -40 | -10 | -12 | -6 |
| Malic acid | Quantifier | 7.4 | 133 | 115 | -50 | -10 | -15 | -12 |
| Malic acid | Qualifier | 7.4 | 133 | 71 | -50 | -10 | -17 | -11 |
| AMP | Quantifier | 7.9 | 346 | 79 | -85 | -10 | -113 | -12 |
| AMP | Qualifier | 7.9 | 346 | 134 | -85 | -10 | -32 | -9 |
| DHAP | Quantifier | 8.2 | 169 | 79 | -31 | -7 | -31 | -7 |
| DHAP | Qualifier | 8.2 | 169 | 97 | -20 | -5 | -31 | -7 |
| Glycerol-3P | Quantifier | 8.2 | 171 | 79 | -20 | -10 | -50 | -9 |
| Glycerol-3P | Qualifier | 8.2 | 171 | 79 | -20 | -10 | -30 | -9 |
| Xy5P | Quantifier | 8.5 | 229 | 79 | -40 | -10 | -59 | -5 |
| Xy5P | Qualifier | 8.5 | 229 | 97 | -40 | -10 | -18 | -5 |
| Xy5P | Qualifier | 8.5 | 229 | 139 | -40 | -10 | -18 | -5 |

| Compound | Type | Retention time | Q1 mass | Q3 mass | DP | EP | CE | CXP |
|---|---|---|---|---|---|---|---|---|
| R5P | Quantifier | 9.0 | 229 | 79 | -40 | -10 | -59 | -5 |
| R5P | Qualifier | 9.0 | 229 | 97 | -40 | -10 | -18 | -5 |
| R5P | Qualifier | 9.0 | 229 | 139 | -40 | -10 | -18 | -5 |
| GAL3P | Quantifier | 9.3 | 169 | 97 | -15 | -10 | -16 | -11 |
| GAL3P | Qualifier | 9.3 | 169 | 79 | -15 | -10 | -16 | -11 |
| E4P | Quantifier | 9.4 | 199 | 97 | -20 | -10 | -20 | -5 |
| E4P | Qualifier | 9.4 | 199 | 79 | -20 | -10 | -20 | -5 |
| ADP-Glc | Quantifier | 9.4 | 588 | 346 | -95 | -10 | -33 | -27 |
| ADP-Glc | Qualifier | 9.4 | 588 | 241 | -95 | -10 | -38 | -30 |
| ADP-Glc | Qualifier | 9.4 | 588 | 134 | -95 | -10 | -89 | -8 |
| UDP-Glc | Quantifier | 10.1 | 565 | 323 | -35 | -10 | -34 | -15 |
| UDP-Glc | Qualifier | 10.1 | 565 | 79 | -80 | -10 | -127 | -4 |
| S7P | Quantifier | 10.3 | 289 | 97 | -35 | -10 | -26 | -9 |
| S7P | Qualifier | 10.3 | 289 | 79 | -35 | -10 | -26 | -9 |
| Glycolate-2P | Quantifier | 10.3 | 155 | 97 | -10 | -10 | -23 | -5 |
| Glycolate-2P | Qualifier | 10.3 | 155 | 81 | -10 | -10 | -23 | -5 |
| PEP | Quantifier | 10.6 | 167 | 79 | -25 | -10 | -23 | -4 |
| PEP | Qualifier | 10.6 | 167 | 63 | -25 | -10 | -92 | -16 |
| 2GPA | Quantifier | 10.4 | 185 | 79 | -30 | -10 | -51 | -4 |
| 2GPA | Qualifier | 10.4 | 185 | 97 | -30 | -10 | -20 | -10 |
| 3PGA | Quantifier | 10.6 | 185 | 79 | -30 | -10 | -51 | -4 |
| 3PGA | Qualifier | 10.6 | 185 | 97 | -30 | -10 | -20 | -10 |
| 3PGA | Qualifier | 10.6 | 185 | 167 | -30 | -10 | -20 | -10 |
| F6P | Quantifier | 9.7 | 259 | 79 | -15 | -10 | -52 | -12 |
| F6P | Quantifier | 9.7 | 259 | 97 | -15 | -10 | -23 | -10 |
| G1P | Quantifier | 10.1 | 259 | 241 | -15 | -10 | -16 | -18 |
| M6P | Quantifier | 10.3 | 259 | 97 | -15 | -10 | -23 | -10 |
| M6P | Qualifier | 10.3 | 259 | 139 | -25 | -10 | -21 | -8 |
| G6P | Quantifier | 10.9 | 259 | 97 | -15 | -10 | -23 | -10 |
| ADP | Quantifier | 10.5 | 426 | 79 | -85 | -10 | -113 | -12 |
| ADP | Qualifier | 10.5 | 426 | 134 | -85 | -10 | -32 | -9 |
| IsoCitric | Quantifier | 10.8 | 191 | 87 | -40 | -10 | -35 | -13 |
| Citric | Quantifier | 10.8 | 191 | 73 | -55 | -10 | -22 | -9 |
| ATP | Quantifier | 12.4 | 506 | 159 | -85 | -10 | -34 | -13 |
| ATP | Qualifier | 12.4 | 506 | 79 | -85 | -10 | -80 | -17 |
| 6-PGluconic Acid | Quantifier | 12.5 | 275 | 97 | -40 | -10 | -21 | -7 |
| 6-PGluconic Acid | Qualifier | 12.5 | 275 | 129 | -40 | -10 | -26 | -8 |
| RuBP | Quantifier | 13.7 | 309 | 79 | -40 | -10 | -64 | -14 |
| RuBP | Qualifier | 13.7 | 309 | 97 | -30 | -10 | -26 | -5 |

| Compound | Type | Retention time | Q1 mass | Q3 mass | DP | EP | CE | CXP |
|---|---|---|---|---|---|---|---|---|
| FBP | Quantifier | 14.0 | 339 | 79 | -35 | -10 | -93 | -2 |
| FBP | Qualifier | 14.0 | 339 | 97 | -35 | -10 | -26 | -7 |
| Sedo-1,7BP | Quantifier | 14.0 | 369 | 97 | -35 | -10 | -75 | -9 |
| Sedo-1,7BP | Qualifier | 14.0 | 369 | 199 | -35 | -10 | -40 | -9 |

*Table S4. Content (μmol $g^{-1}$ Fresh Weight) of metabolites found in Arabidopsis thaliana (n=3) under effect of pestle and cooling*

| Compound | Category | No Pestle, No cooling | Pestle, No cooling | No Pestle, Cooling | Pestle, Cooling |
|---|---|---|---|---|---|
| Trp | Amino Acid | 26.6 ± 4.4 | 22.0 ± 0.5 | 20.3 ± 1.3 | 21.7 ± 1.8 |
| Phe | Amino Acid | 77.8 ± 10.8 | 69.1 ± 1.6 | 60.7 ± 3.3 | 67.4 ± 4.6 |
| ILe | Amino Acid | 59.1 ± 10.2 | 51.5 ± 1.2 | 43.0 ± 1.3 | 52.2 ± 4.5 |
| Leu | Amino Acid | 65.2 ± 10.2 | 55.9 ± 0.8 | 47.8 ± 2.8 | 55.5 ± 4.4 |
| Met | Amino Acid | 14.7 ± 1.8 | 12.6 ± 1.5 | 12.4 ± 0.8 | 13.0 ± 1.1 |
| Val | Amino Acid | 174.8 ± 29.0 | 136.3 ± 8.6 | 122.7 ± 15.3 | 129.4 ± 22.5 |
| Tyr | Amino Acid | 25.2 ± 3.4 | 21.5 ± 1.3 | 18.3 ± 1.4 | 19.0 ± 1.2 |
| Pro[a] | Amino Acid | 1073.0 ± 62.7 | 1014.5 ± 77.2 | 972.1 ± 73.8 | 949.3 ± 44.9 |
| HPro | Amino Acid | 8.6 ± 0.8 | 6.8 ± 0.8 | 6.8 ± 0.4 | 6.8 ± 0.1 |
| Ala[a] | Amino Acid | 683.8 ± 44.4 | 651.8 ± 67.4 | 652.0 ± 68.6 | 613.1 ± 6.8 |
| Thr[a] | Amino Acid | 1102.7 ± 109.1 | 1022.7 ± 80.4 | 1031.5 ± 77.4 | 957.1 ± 20.7 |
| Gly | Amino Acid | 205.5 ± 18.4 | 184.7 ± 13.8 | 210.3 ± 22.2 | 166.4 ± 9.7 |
| Gln[a] | Amino Acid | 3078.1 ± 224.3 | 2883.8 ± 243.0 | 2892.8 ± 229.4 | 2737.8 ± 27.4 |
| Ser[a] | Amino Acid | 1698.0 ± 133.4 | 1565.8 ± 150.0 | 1540.7 ± 119.3 | 1443.0 ± 23.7 |
| Asn[a] | Amino Acid | 801.0 ± 76.8 | 747.4 ± 77.5 | 768.8 ± 64.4 | 706.7 ± 16.7 |
| Glu[b] | Amino Acid | 19509.4 ± 1264.9 | 18121.5 ± 2172.1 | 18257.6 ± 1502.9 | 15727.5 ± 683.3 |
| His | Amino Acid | 152.5 ± 24.3 | 134.6 ± 6.8 | 124.8 ± 9.2 | 125.0 ± 3.5 |
| Asp[a] | Amino Acid | 2882.1 ± 288.2 | 2542.5 ± 338.0 | 2515.2 ± 187.9 | 2270.3 ± 160.8 |
| Arg | Amino Acid | 76.5 ± 3.8 | 77.1 ± 6.2 | 64.9 ± 13.4 | 67.6 ± 2.0 |
| Orn | Amino Acid | 8.0 ± 1.8 | 6.6 ± 1.1 | 6.5 ± 2.1 | 5.2 ± 0.8 |
| Lys | Amino Acid | 69.6 ± 15.7 | 53.6 ± 2.3 | 48.2 ± 8.3 | 49.3 ± 3.7 |
| Acetyl-COA | Coenzyme | ND | ND | ND | ND |
| AMP | Nucleotide / Phosphate | 10.5 ± 2.7 | 7.5 ± 2.5 | 9.7 ± 1.9 | 7.9 ± 1.1 |
| ADP | Nucleotide / Phosphate | 11.9 ± 3.3 | 9.4 ± 2.2 | 8.6 ± 1.5 | 8.0 ± 2.9 |
| ATP | Nucleotide / Phosphate | 40.5 ± 8.8 | 28.0 ± 8.0 | 24.0 ± 4.2 | 23.7 ± 8.0 |

| Compound | Category | No Pestle, No cooling | Pestle, No cooling | No Pestle, Cooling | Pestle, Cooling |
|---|---|---|---|---|---|
| ADP-Glc | Nucleotide Sugar | 1.2 ± 0.2 | 0.9 ± 0.1 | 0.8 ± 0.0 | 0.8 ± 0.1 |
| UDP-Glc | Nucleotide Sugar | 92.1 ± 21.7 | 61.2 ± 6.2 | 62.5 ± 7.9 | 57.7 ± 6.6 |
| Ascorbic acid | Organic Acid | 3082.3 ± 6.7 | 3062.1 ± 18.1 | 3117.1 ± 115.2 | 3033.8 ± 8.9 |
| Pyruvic acid | Organic Acid | 40.8 ± 6.4 | 38.3 ± 4.6 | 37.9 ± 0.9 | 33.0 ± 1.4 |
| Glycolic acid | Organic Acid | 46.8 ± 3.2 | 45.6 ± 6.9 | 40.5 ± 4.1 | 39.6 ± 4.5 |
| Glyceric acid | Organic Acid | 162.8 ± 42.3 | 127.6 ± 35.3 | 133.6 ± 16.3 | 104.4 ± 25.3 |
| Shikimic acid | Organic Acid | 31.7 ± 2.1 | 29.2 ± 3.4 | 27.7 ± 3.3 | 25.2 ± 0.6 |
| 2-Ketoglutaric acid[a] | Organic Acid | 368.3 ± 38.3 | 325.7 ± 13.8 | 332.2 ± 42.5 | 328.3 ± 7.0 |
| Succinic acid | Organic Acid | 332.4 ± 15.9 | 326.4 ± 23.6 | 304.2 ± 41.4 | 274.7 ± 5.6 |
| Fumaric acid[b] | Organic Acid | 9002.3 ± 613.7 | 8627.5 ± 799.8 | 7754.7 ± 491.6 | 7049.2 ± 391.0 |
| Malic acid[b] | Organic Acid | 3546.0 ± 168.9 | 3257.6 ± 339.7 | 3016.0 ± 129.4 | 2872.2 ± 60.5 |
| Glyoxylic acid | Organic Acid | 63.0 ± 2.9 | 63.4 ± 3.0 | 65.2 ± 0.2 | 59.1 ± 2.3 |
| Citric acid[b] | Organic Acid | 8837.1 ± 1328.8 | 8369.1 ± 2251.1 | 7597.9 ± 496.3 | 6439.1 ± 525.3 |
| Isocitric acid | Organic Acid | 233.5 ± 40.4 | 186.0 ± 33.0 | 178.3 ± 2.9 | 169.6 ± 12.9 |
| E4P | Phosphorylated Metabolite | ND | ND | ND | ND |
| Glycolate-2P | Phosphorylated Metabolite | ND | ND | ND | ND |
| DHAP | Phosphorylated Metabolite | 57.9 ± 4.7 | 52.8 ± 4.1 | 50.7 ± 9.0 | 47.5 ± 3.6 |
| Glycerol-3P | Phosphorylated Metabolite | 46.2 ± 17.5 | 34.8 ± 8.8 | 29.5 ± 6.1 | 28.5 ± 9.0 |
| Xy5P+Ru5P[1] | Phosphorylated Metabolite | 55.8 ± 16.7 | 21.0 ± 13.9 | 36.3 ± 17.4 | 19.4 ± 17.8 |
| GAL3P | Phosphorylated Metabolite | 23.0 ± 1.2 | 21.1 ± 0.6 | 19.6 ± 0.3 | 19.5 ± 1.1 |
| S7P | Phosphorylated Metabolite | 21.0 ± 3.7 | 17.7 ± 1.2 | 16.6 ± 1.4 | 16.5 ± 1.7 |
| 6-PGluconic Acid | Phosphorylated Metabolite | 1.4 ± 0.1 | 1.1 ± 0.2 | 1.0 ± 0.1 | 1.0 ± 0.1 |
| RuBP | Phosphorylated Metabolite | 46.8 ± 19.7 | 42.1 ± 7.4 | 41.5 ± 23.1 | 47.0 ± 15.2 |
| FBP | Phosphorylated Metabolite | 19.9 ± 1.8 | 15.9 ± 3.5 | 15.0 ± 2.0 | 15.5 ± 1.2 |
| F6P | Phosphorylated Metabolite | 83.9 ± 24.2 | 53.3 ± 6.1 | 52.2 ± 6.0 | 48.3 ± 9.4 |
| G1P | Phosphorylated Metabolite | 27.0 ± 8.1 | 15.4 ± 2.9 | 16.2 ± 1.1 | 15.0 ± 3.8 |

| Compound | Category | No Pestle, No cooling | Pestle, No cooling | No Pestle, Cooling | Pestle, Cooling |
|---|---|---|---|---|---|
| M6P | Phosphorylated Metabolite | 17.6 ± 3.2 | 14.8 ± 1.6 | 14.4 ± 2.4 | 13.0 ± 1.1 |
| G6P | Phosphorylated Metabolite | 42.2 ± 3.6 | 42.6 ± 2.3 | 39.3 ± 7.6 | 37.7 ± 3.0 |
| PEP | Phosphorylated Metabolite | 2.1 ± 0.2 | 1.7 ± 1.0 | 1.3 ± 0.6 | 1.4 ± 0.3 |
| 2PGA | Phosphorylated Metabolite | 4.1 ± 0.6 | 3.3 ± 0.3 | 3.3 ± 0.1 | 3.3 ± 0.3 |
| 3PGA | Phosphorylated Metabolite | 43.7 ± 9.7 | 43.2 ± 11.0 | 38.9 ± 10.8 | 37.7 ± 4.0 |
| Fructose[2] | Sugar | 239.8 ± 48.0 | 245.3 ± 90.6 | 221.0 ± 42.8 | 156.2 ± 21.5 |
| Glucose[2] | Sugar | 1557.6 ± 49.8 | 1623.5 ± 164.9 | 1528.2 ± 28.7 | 1361.7 ± 36.4 |
| Sucrose[a] | Sugar | 1177.0 ± 141.7 | 1079.6 ± 77.3 | 1128.2 ± 133.4 | 1147.8 ± 23.3 |
| Trehalose | Sugar | 13.2 ± 5.1 | 10.0 ± 4.0 | 8.0 ± 1.4 | 8.1 ± 0.4 |

*ND: Not detected, NA: Not applicable*

*[a] quantify from diluted 20-fold extractant, [b] quantify from diluted 50-fold extractant*

*[1] Xy5P+Ru5P were quantified together based on Xy5P standard.*

*[2] Fructose and Glucose were measured with ion source temperature at 200°C*

*MDP concentration optimization*

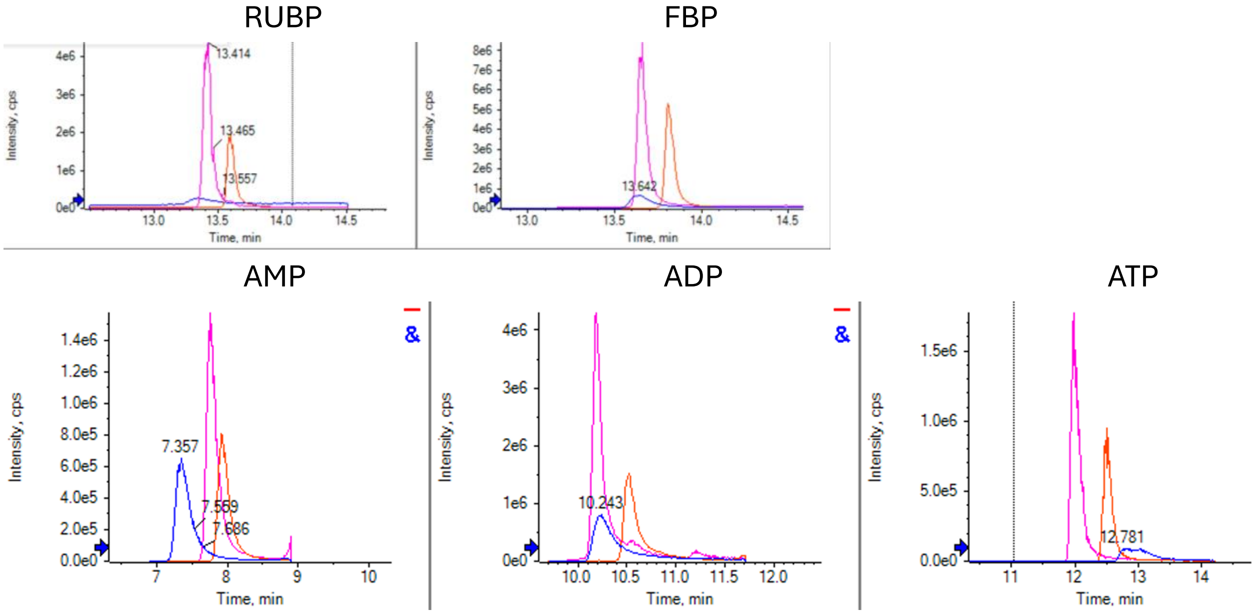


*Figure S1. Representative chromatograms of phosphorylated metabolites using mobile phase A containing 10 μM MDP, 20μM MDP, 20 μM MDP with 0.01% Formic acid.*

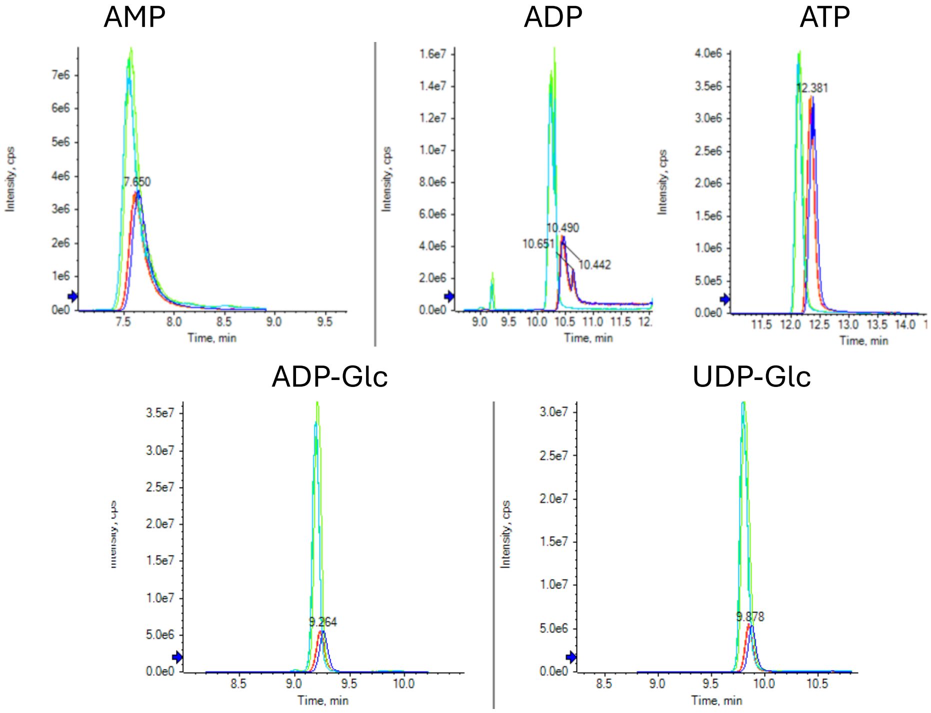


*Figure S2. Representative chromatograms of phosphorylated metabolites using mobile phase A containing 50 μM MDP and 75 μM MDP*

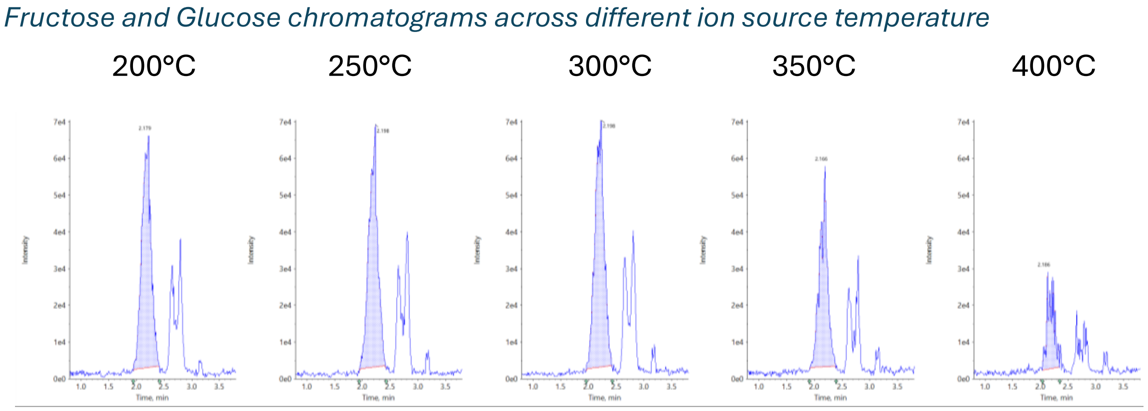


*Figure S3. Fructose and Glucose chromatograms across different ion source temperatures.*

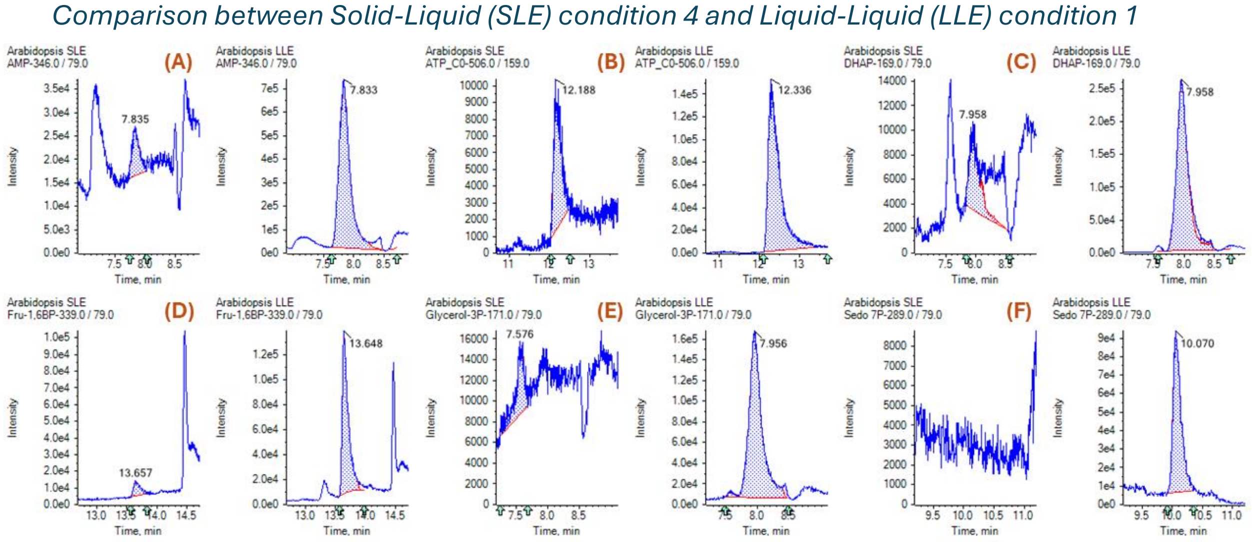


*Figure S4. Representative chromatographic comparison between Solid-Liquid (SLE) condition 4 and Liquid-Liquid (LLE) condition 1 of (A): AMP, (B): ATP, ((C): DHAP, (D): FBP, (E): Glycerol-3P, (F): S7P*
